# Weak and inverse latitudinal diversity gradients in the globally dominant flying insect clades

**DOI:** 10.1101/2025.04.29.651164

**Authors:** Bernardo F. Santos, Amrita Srivathsan, Karen Neves, Rudolf Meier

## Abstract

Latitudinal diversity gradients are a key concept in biodiversity science, but the available data are skewed towards large-bodied, charismatic taxa. To address this bias, we analyzed DNA barcode data for 1.35 million specimens of flying insects collected using 101 Malaise traps in 27 countries across six continents. We identify those ten families that consistently contribute more than half of all specimens and species across climates, habitats, and continents. Contrary to canonical expectations, five families have inverse or non-significant latitudinal gradients while the remaining five exhibit only weak to average gradients compared to 470 other terrestrial datasets. Furthermore, precipitation seasonality, rather than temperature, emerged as the strongest climatic correlate. Overall, we thus not only reveal a surprising convergence in the globally dominant insect clades but also illustrate that key textbook concepts in macroecology need testing for poorly studied invertebrate taxa that contribute most of Earth’s animal biodiversity and may behave unexpectedly.

## Introduction

A central expectation in biodiversity science is that for most clades species richness increases toward the equator, a pattern known as the latitudinal diversity gradient (Darwin 1859; Pianka 1966; Hawkins 2001; Hillebrand 2004). Although numerous ecological and evolutionary explanations have been proposed to account for this pattern, no clear consensus has emerged (Pianka 1966; Fine 2015). Ecological hypotheses emphasize the influence of environmental factors such as rainfall and primary productivity, whereas evolutionary perspectives focus on differences in diversification and extinction rates, along with systematic disparities in the time available for diversification between tropical and temperate regions (Willig et al. 2003; Mittelbach et al. 2007). Effectively, some imply that tropical environments support higher species richness even at local levels (alpha diversity) through mechanisms such as elevated energy availability (Currie et al. 2004), greater climatic stability (Jetz & Fine 2012; Fine 2015) or intensified biotic interactions that enable finer niche partitioning (Schemske 2009). Others emphasize processes that could drive greater species turnover across space (beta diversity), such as environmental heterogeneity (Stein et al. 2014), dispersal limitation (Morlon et al. 2011), or historical patterns of speciation and extinction (Mittelbach et al. 2007; Wiens & Donoghue 2004). It may thus not surprise that a recent synthesis (Kinlock et al. 2018) of the empirical evidence from 970 datasets found broadly consistent LDG patterns for alpha and beta diversity estimates based on similar numbers of studies.

Despite the importance of the LDG in ecological and evolutionary theory, most empirical studies focus on well-known taxa such as vertebrates, flowering plants, and other charismatic organisms. In contrast, many species-rich invertebrate clades, which contain most multicellular species, have received little attention. For example, among the 470 terrestrial studies included in the meta-analyses of Hillebrand (2004) and Kinlock et al. (2018), more than 60% are vertebrates and vascular plants. Invertebrate data, when present, tend to be for taxa with relatively large species such as bees, butterflies, and dragonflies. This bias likely reflects the difficulty of compiling species lists or obtaining distribution data for small invertebrate taxa that tend to be data-deficient, even though they comprise most species and a large share of global biomass (Rohr et al. 2007; Wilson 2017). As a result, biodiversity patterns for small species remain comparatively poorly understood although there is tentative evidence that they may not always follow the same trends observed in larger organisms (Hillebrand and Azovsky 2001; Hillebrand 2004; Kinlock et al. 2018).

Of particular concern is that data for latitudinal diversity gradients are almost entirely lacking for so-called “dark taxa,” defined as clades in which fewer than 10 % of species have been described and total diversity exceeds 1,000 species (Hartop et al. 2022). These taxa frequently dominate biodiversity samples collected with widely used biomonitoring tools such as Malaise and pitfall traps, yet dark taxa remain largely absent from global biodiversity analyses. The primary obstacle is taxonomic neglect (Srivathsan et al. 2023), which gives rise to two well-known shortfalls: the Linnean shortfall, referring to the large number of species that remain undescribed, and the Wallacean shortfall, referring to the lack of geographic data for species (Lomolino 2004). For dark taxa, the Linnean shortfall is especially severe and contributes to an even deeper Wallacean shortfall. Indeed, two self-reinforcing dynamics are at play. First, the lack of taxonomic and distributional data makes dark taxa less likely to be included in broad-scale biodiversity studies. Second, Srivathsan et al. (2023) show that taxonomic neglect increases with species diversity, and that most flying insect species belong to clades dominated by small species. As a result, the taxa that contribute the largest number of species to global biodiversity are also those most affected by the Linnean and Wallacean shortfalls, which limits the development of distributional data.

Recent methodological advances now offer a path forward, enabling standardized, spatially explicit sampling of dark taxa at global scales. By combining tools such as Malaise and pitfall traps (Noyes 1989a; Karlsson et al. 2020; Skvarla et al. 2021) with large-scale DNA barcoding of individual specimens (“megabarcoding”: Chua et al. 2023), it is now possible to generate species-level data suitable for global comparisons. Barcodes can be used to assign specimens to putative species, a procedure that is sufficiently precise to support comparative studies of species-level diversity patterns of invertebrate communities (Telfer et al. 2015; Geiger et al. 2016; Ashfaq et al. 2017; DeWaard et al. 2019a; D’Souza et al. 2021; Yeo et al. 2021; Chimeno et al. 2023; Srivathsan et al. 2023; Seymour et al. 2024). This approach can yield important data for sites that span broad latitudinal gradients as well as a range of continents and habitats, allowing for the analysis of taxonomic composition and the testing of textbook generalizations about global biodiversity patterns..

Analyses of flying insect diversity at a global scale are starting to yield both surprising and expected insights. One study found that most species in flying insect communities belong to a small number of hyperdiverse families, based on Malaise trap samples from eight countries on five continents (Srivathsan et al. 2023). The same ten families accounted for roughly half of all species and specimens across continents, climates, and habitat types. Another study using data from 129 sites in the Global Malaise Trap Project (Seymour et al. 2024) found evidence for a higher species turnover at lower latitudes. What is more surprising is that these two studies are only among the first to describe global patterns for the flying insects commonly sampled with Malaise traps, the most widely used method for insect biomonitoring. The paucity of such studies documents how little we know about these samples despite their central role in monitoring the most species-rich animal taxon.

We here use data from 101 Malaise traps placed at 45 localities in 27 countries across six continents and all biogeographic regions to carry out a global test of Srivathsan et al.’s (2023) proposal that ten dominant flying insect clades contribute around 50 % of all species level diversity in Malaise trap samples. We also analyze latitudinal diversity gradients for each of these taxa and examine whether the same clades dominate when only tropical sites are considered. As part of this broader analysis, we pay particular attention to ichneumonid wasps, for which an inverse latitudinal diversity gradient had long been proposed (Owen and Owen 1974; Janzen and Pond 1975; Janzen 1981; Gauld 1986; Sime and Brower 1998; Castellanos-Labarcena et al. 2024). However, the available evidence had largely come from studies based on a small number of sites (Owen and Owen 1974; Janzen and Pond 1975; Noyes 1989b), geographically restricted sampling (Janzen 1981; Gauld 1986), data derived from catalogs and checklists, or samples that were not standardized prior to analysis (Castellanos-Labarcena et al. 2024). To address these shortcomings, we can here use standardized samples and include other parasitoid clades in the analysis to evaluate whether an inverse gradient in ichneumonid diversity exists and whether it can be explained by parasitoidism or host choice.

## Results

### A Few Insect Families Dominate Global Flying Insect Diversity

We analyzed data from 101 Malaise trap samples comprising 1,363,772 specimens, of which 1,280,387 were identified as insects. Datasets were obtained either from published studies or public projects at the Barcode of Life Data System (Supplementary Material S1). Species diversity was estimated using a 3% sequence dissimilarity threshold applied to DNA barcodes, yielding between 56 and 8,223 molecular operational taxonomic units (mOTUs) per trap (mean 2,089; standard deviation 1,772; Supplementary Material S1). We estimate the total number of species in the study to likely exceed 130,000, given the high diversity within the most common families and the low overlap in species composition across sites. For the top ten insect families alone, we recorded more than 66,000 species-level mOTUs and species turnover was high: only 12.6 % of mOTUs were found at more than one site (range 8.54 to 18.2 %), and fewer than 0.2 % occurred in ten or more sites (Supplementary Material S3).

Faunal analysis at species level confirms the dominance of a small number of highly diverse and abundant insect families. The ten most diverse families account on average for 52.4% of species and 59.7 % of specimens in each sample (Table 1; Fig. S1; Supplementary Material S2). All ten families belong to just three insect orders: Diptera (six families: Cecidomyiidae, Chironomidae, Sciaridae, Phoridae, Ceratopogonidae, and Mycetophilidae), Hymenoptera (three families: Ichneumonidae, Platygastridae, and Braconidae), and Hemiptera (Cicadellidae). Families ranked 11th to 20th in diversity contributed an additional 11.3 % of species and 13.7 % of specimens, while the remaining 36.3% of species were distributed across the other 643 families. Eight of the global top-ten families also ranked among the top-ten when only tropical sites were considered. The only changes were the replacement of Ichneumonidae (ranked 4th globally) and Mycetophilidae (10th) by Formicidae (global rank 12th) and Psychodidae (global rank 14th) in the tropical analysis.

**Table 1.** The top 10 most diverse families make up over 50% of the specimens and species found in the catch of Malaise trap samples across the world.

| Order | Family (average size) | # specimens | % specimens | # species | % species |
| --- | --- | --- | --- | --- | --- |
| Diptera | Cecidomyiidae (2.8 mm) | 175,668 | 12.24% | 26,387 | 16.16% |
| Diptera | Chironomidae (3.6 mm) | 146,323 | 15.97% | 5,177 | 7.79% |
| Diptera | Sciaridae (3.3 mm) | 126,731 | 8.90% | 5,315 | 4.78% |
| Hymenoptera | Ichneumonidae (10.9 mm) | 21,354 | 2.05% | 4,563 | 4.63% |
| Hymenoptera | Platygasteridae (1.6 mm) | 39,679 | 3.09% | 6,200 | 4.43% |
| Diptera | Phoridae (1.7 mm) | 65,407 | 4.85% | 5,473 | 3.95% |
| Hymenoptera | Braconidae (5.5 mm) | 19,161 | 1.61% | 4,634 | 3.46% |
| Diptera | Ceratopogonidae (2.4 mm) | 73,656 | 5.54% | 4,080 | 3.11% |
| Hemiptera | Cicadellidae (6.9 mm) | 45,545 | 3.86% | 2,867 | 2.15% |
| Diptera | Mycetophilidae (5.4 mm) | 18,069 | 1.60% | 1,809 | 1.91% |
| **Total** |  | **731,593** | **59.71%** | **66,505** | **52.38%** |

When families were ranked by abundance rather than species diversity, the ten most abundant families contributed on average 62.7 % of all specimens per sample. The taxonomic composition of these top-ten based on abundance changed only slightly compared to the composition of the top-ten based on diversity, with Braconidae (ranked 6th by species) and Mycetophilidae (10th by species) replaced by Formicidae and Psychodidae.

### The top 10 taxa have weak latitudinal diversity gradients

We established the latitudinal diversity gradients for the top-ten most diverse insect families (Fig. 1) using both species richness and Shannon’s diversity index (Magurran and McGill 2011), with diversity measures adjusted for sample size (Supplementary Material S3). We then compared the results to established gradients in 470 terrestrial taxa compiled in two comprehensive meta-analyses (Hillebrand 2004; Kinlock et al. 2018). Effect sizes were calculated as Pearson’s correlation coefficient (r) between species richness and latitude, then normalized using Fisher’s z transformation, following the methods of Hillebrand (2004) and Kinlock et al. (2018).

**Figure 1.**
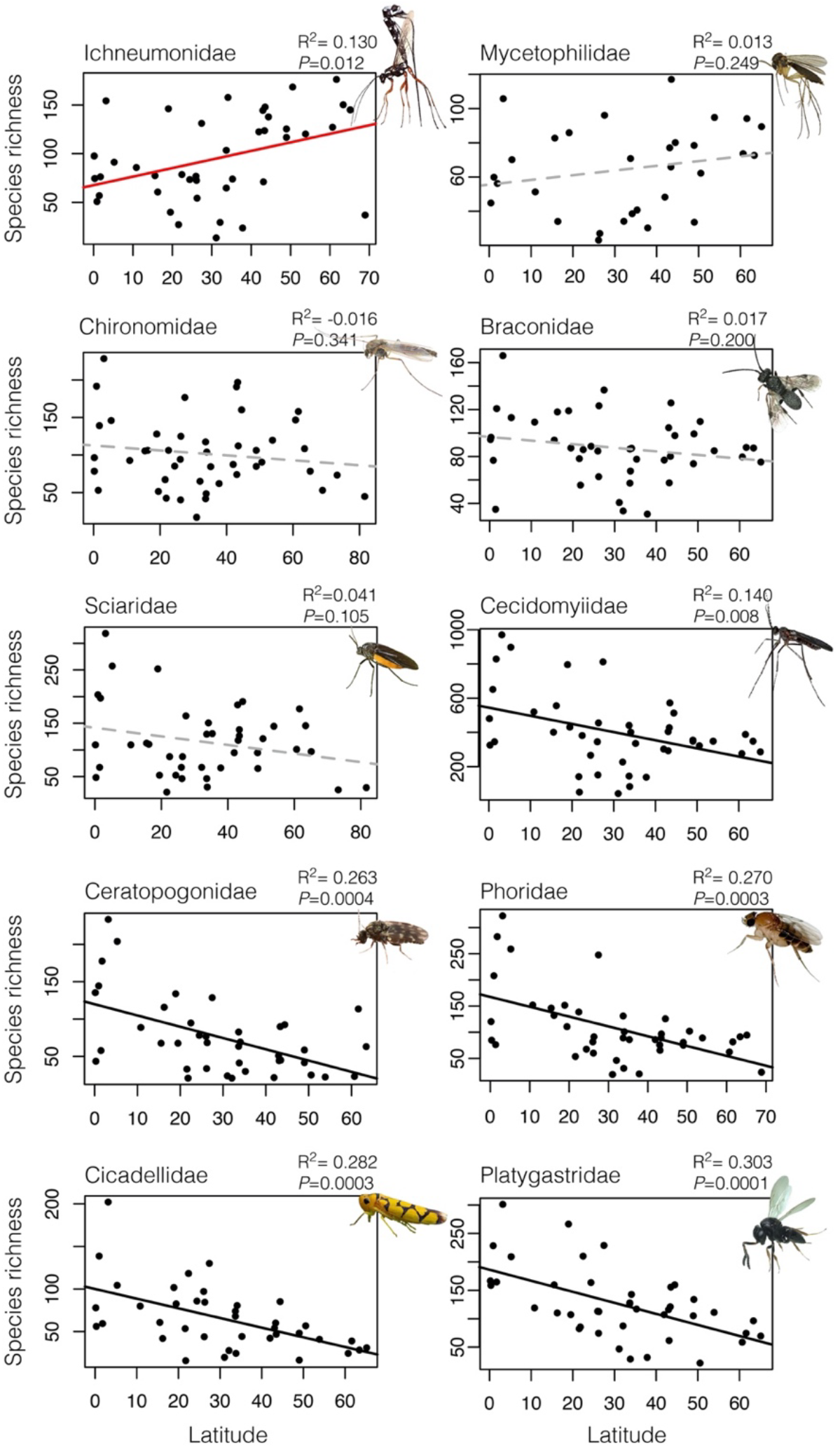
The top 10 insect families show unexpected variation in latitudinal diversity. Linear regression of normalized species richness against latitude for the top ten families. Red trendline is for the only significant inverse gradient, dashed gray for non-significant gradients (four families), and black for significant positive gradients (six families).

Contrary to expectations, we find that latitudinal diversity gradients are comparatively weak, absent, or even reversed. Indeed, half of the ten most diverse insect families show either a non-significant pattern (Braconidae, Chironomidae, Mycetophilidae, Sciaridae; R^2^ < 0.04, P > 0.10, n = 31–45) or an inverse gradient (Ichneumonidae: R^2^ = 0.1298, P = 0.0119, n = 41). The remaining five families (Cecidomyiidae, Ceratopogonidae, Cicadellidae, Phoridae, Platygastridae) exhibit only weak to average gradients, with effect sizes clustering around the 50th percentile of gradient strength observed across 470 terrestrial taxa in published meta-analyses (Figs 2, S2; Supplementary Material S5). To test whether this lack of strong LDGs could be due to chance, we drew 10,000 random samples of 10 LDGs from the literature sample and compared the strongest from each iteration to our strongest (Platygastridae, R^2^ = –0.642). The probability of obtaining no stronger effect than ours in such samples was extremely low (*P* = 0.0008), a result that remained significant even when restricted to alpha-diversity studies (*P* = 0.0066). A Wilcoxon rank sum test furthermore confirmed that our effect sizes are skewed toward the weak end of the LDG distribution (W = 3211, *P* = 0.0196), although this was not significant for alpha-diversity data alone (W = 1188, *P* = 0.1034).

**Figure 2.**
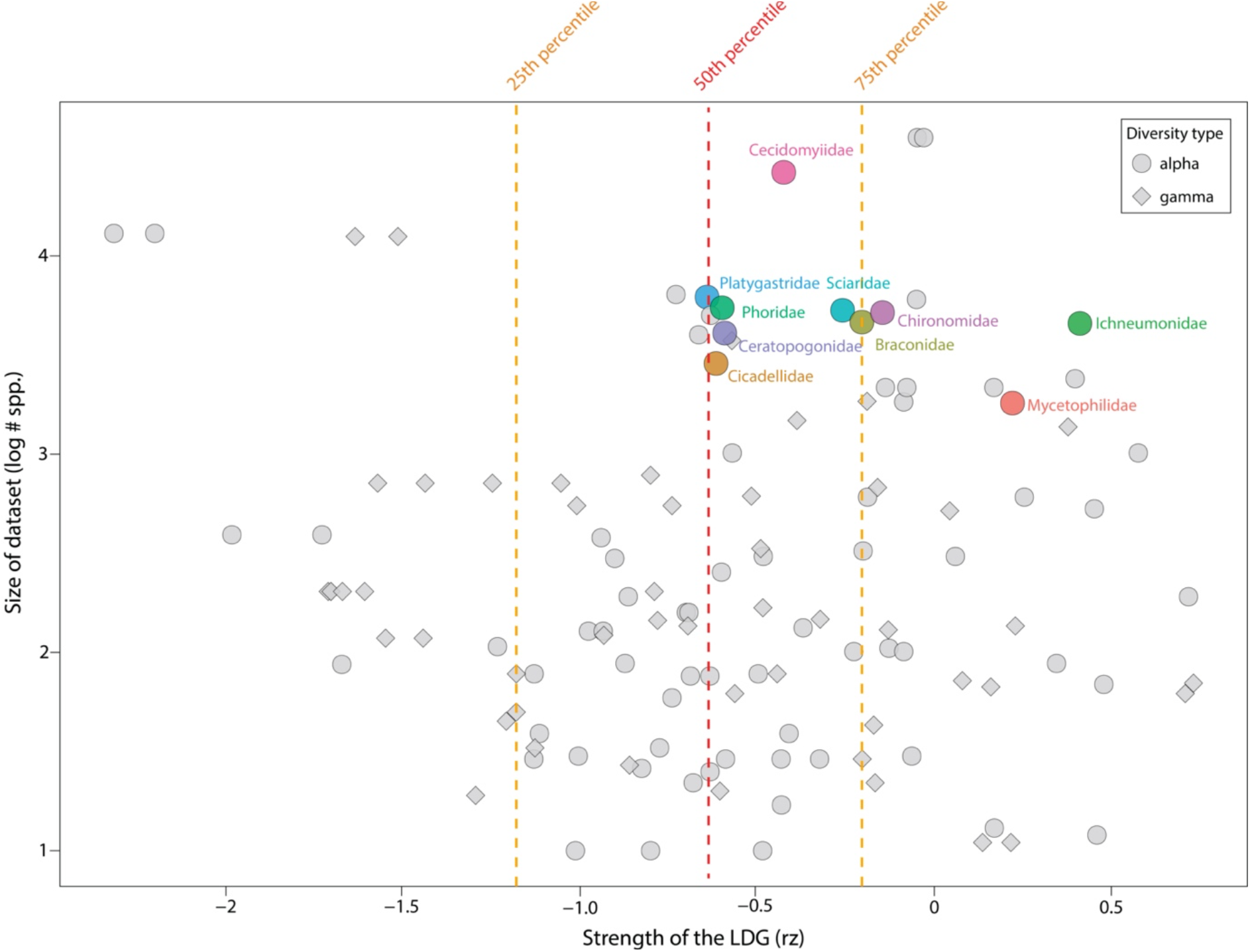
Most of the top 10 insect families have below-average latitudinal diversity gradients based on large and well-sampled datasets. Z-transformed correlation coefficients indicate gradient strength, with percentile lines based on 470 terrestrial datasets from Hillebrand (2004) and Kinlock et al. (2018). Colored dots represent the top 10 families from this study; gray dots represent 114 datasets with known species richness.

When all species in the samples are analyzed simultaneously (Figs 3, S3), a significant albeit weak latitudinal gradient is observed further suggesting that the top-ten families have unusual gradients. For the full dataset, species diversity increases by an average of 1.27 % for each degree of latitude away from the equator, although substantial variance remains within the same latitudinal bands (R^2^ = 0.23, P = 0.0005, n = 45).

**Figure 3.**
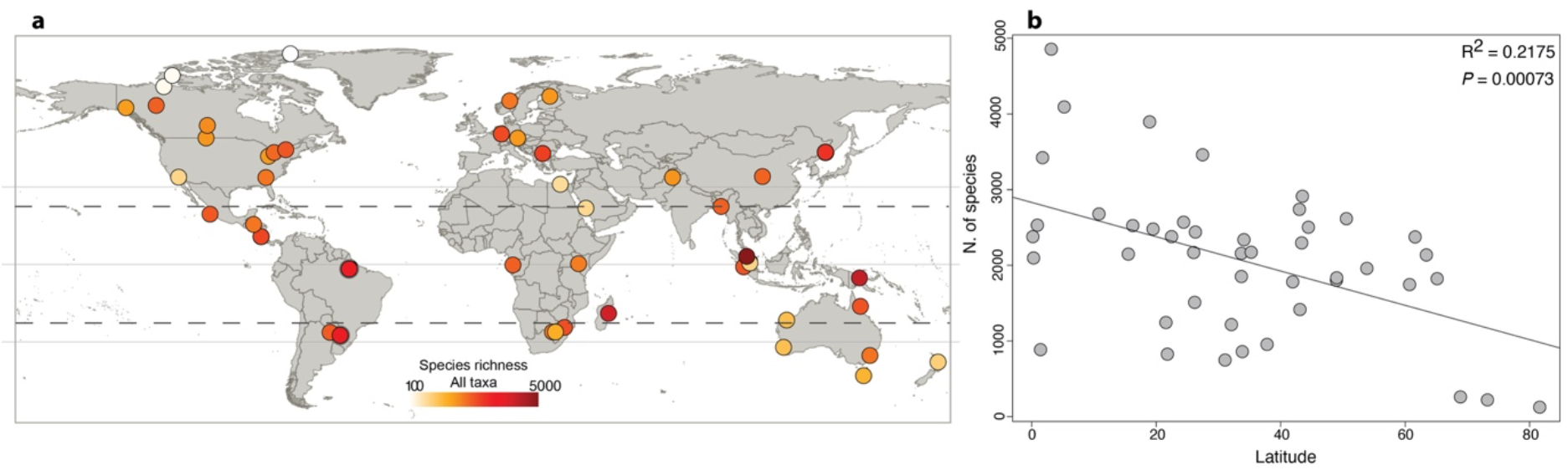
A significant but weak latitudinal diversity gradient is observed for all flying insect species represented in the global sample. a, Map showing the location of the 45 trapping sites, with colors indicating standardized species richness estimates based on 13,500 specimens per site. b, weak but significant latitudinal diversity gradient when species richness is regressed against latitude.

To explore potential drivers of the observed diversity patterns, we used multi-model averaging (Bartoń 2023; Naimi et al. 2014) to identify bioclimatic variables associated with global diversity in each of the top ten families and in the aggregate dataset. The best supported model, which included six temperature and precipitation variables, explained 33 to 53 % of the variation in global diversity based on adjusted R^2^ values (Supplementary Material S4). Overall, precipitation-related variables showed stronger correlations than temperature variables, with precipitation seasonality ranking highest in weighted AIC and showing the most frequent significant associations across taxa (Fig. S4). Most taxa exhibited broadly similar responses to the climatic variables. However, Ichneumonidae stood out as the only family with a significant positive association with temperature seasonality. Other families showing weak or inverse latitudinal gradients, such as Mycetophilidae and Chironomidae, also tended to respond positively to increased temperature seasonality (Fig. S4; Supplementary Material S4).

### An inverse latitudinal gradient for Ichneumonidae (Hymenoptera)

A significant inverse latitudinal diversity gradient was observed in one of the top-ten taxa, ichneumonid wasps. The effect size observed for Ichneumonidae places them near the fifth percentile among the most extreme inverse gradients reported in the literature (Figs 2, S2). The result was stable regardless of whether species richness or Shannon’s diversity index was used to measure diversity. It also remained robust when sample size is normalized using either sample coverage (Roswell et al. 2021) or specimen level rarefaction (Supplementary Material S3).

This inverse latitudinal diversity gradient is partly driven by increasing relative abundance at higher latitudes. As a proportion of the total catch in each trap, ichneumonids increase from less than 1% of specimens near the equator to over 5% in most samples between 50° and 65° latitude (R^2^ = 0.51, P = 5.593 × 10^−8^, n = 43; Fig. 4; Supplementary Material S1). The proportional species richness of Ichneumonidae also increases toward temperate regions, reaching more than 10 % of all captured species in some northern temperate samples (R^2^ = 0.67, P = 1.841 × 10^−11^, n = 43; Fig. 4). This effect is more pronounced in the Northern (abundance R^2^ = 0.56; diversity R^2^ = 0.73) than the Southern Hemisphere (abundance R^2^ = 0.15; diversity R^2^ = 0.34). Beyond the Arctic Circle, both the relative abundance and diversity of Ichneumonidae decline sharply, which somewhat weakens the overall inverse latitudinal gradient when Arctic samples are included. Thus, after excluding the Arctic sites, the inverse diversity gradient is even stronger (R^2^ = 0.217, P = 0.0014) highlighting temperate regions as the main diversity hotspot for this family.

**Figure 4.**
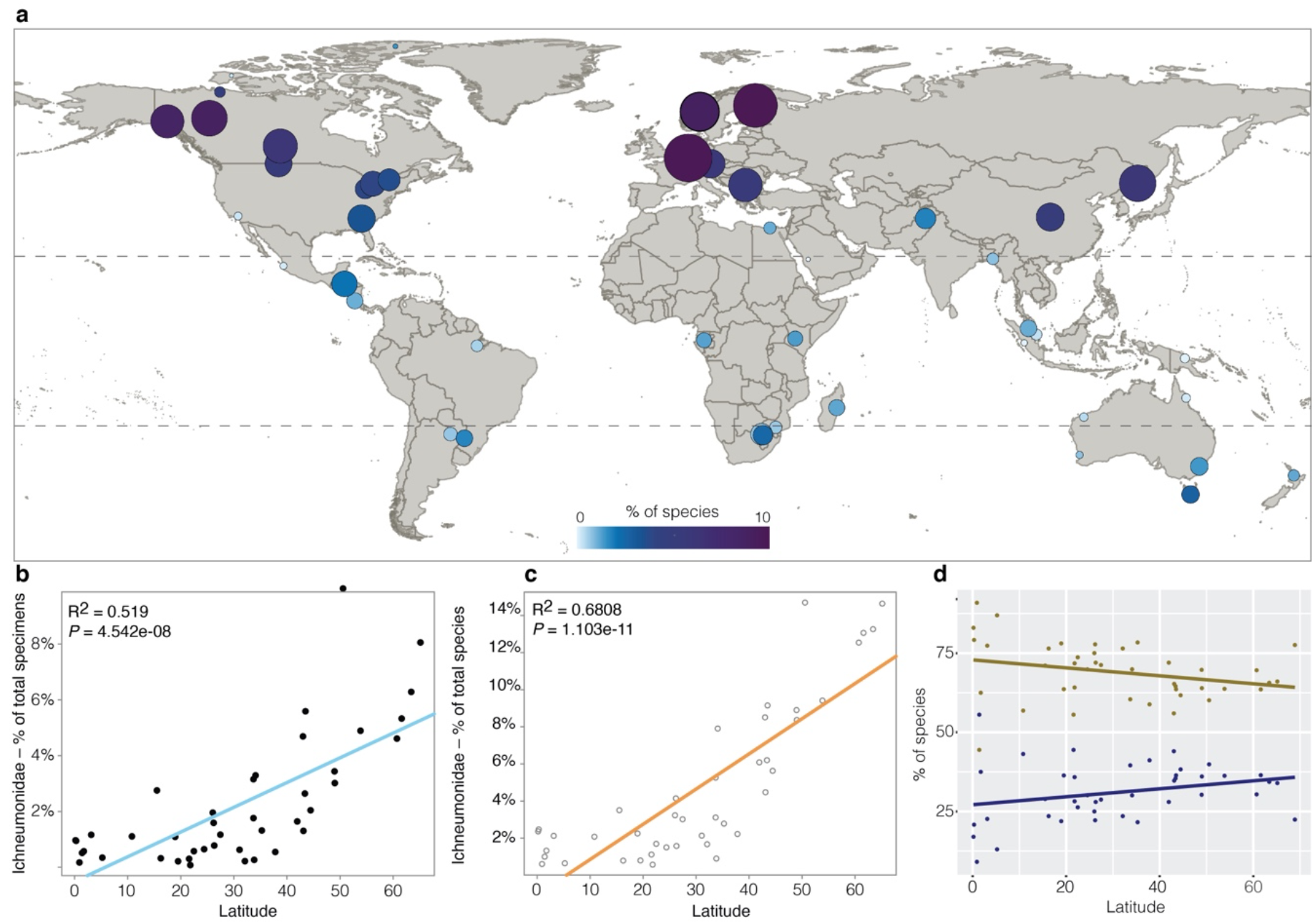
Ichneumonidae are most diverse and abundant in temperate zones. a, Geographic distribution of Ichneumidae as a proportion of total specimens (dot size) and species (color gradient) across sampled sites. b, Linear regression of ichneumonid specimen proportion against latitude. c, Linear regression of ichneumonid species proportion against latitude. d, Proportion of specialist (koinobiont) and generalist (idiobiont) ichneumonid species across sites.

To test whether the inverse latitudinal diversity gradient in Ichneumonidae was related to differences in host use, we assigned mOTUs to subfamilies using BLAST. Each subfamily was then classified as either idiobiont (“generalist”) or koinobiont (“specialist”; see definitions in Askew and Shaw 1986). We calculated the proportion of koinobiont and idiobiont species in each sample and computed site-specific diversity metrics for both groups separately. The proportion of koinobiont species increased slightly toward the tropics, but the relationship was not statistically significant (R^2^ = 0.049, P = 0.087, n = 45; Fig. 4d, Supplementary Material S6). Both groups showed increasing diversity toward the temperate zone, with nearly parallel regression lines (Fig. S5). Removing two subfamilies largely restricted to temperate zones (Ctenopelmatinae and Diplazontinae) had only a minor effect on the overall gradient (R^2^ = 0.124, P = 0.024, n = 41; Supplementary Material S7), suggesting that the inverse latitudinal diversity gradient is a general feature of Ichneumonidae rather than being driven by specific subfamilies or host-use strategies.

## Discussion

Our understanding of global biodiversity remains shaped by a narrow subset of species, biased toward large, well-known clades of animals, and plants that are often comparatively species-poor. Latitudinal diversity gradients are no exception. Despite their central role in ecological theory, studies of these gradients have traditionally neglected many hyperdiverse invertebrate taxa. For example, among the 470 terrestrial datasets included in recent meta-analyses (Hillebrand 2004; Kinlock et al. 2018), only about one fifth focus on insects, and most of those involve large-bodied groups such as bees, butterflies, and dragonflies. Diptera, which are here revealed to include six of the ten most diverse families of flying insects, are entirely absent. Indeed, only a few high-diversity insect clades, such as bees, ants and ichneumonid wasps, have ever been included (e.g. Owen & Owen 1974; Economo et al. 2018).

The scarcity of global diversity studies targeting the most speciose taxa is unsusprising when considering that, until recently, we lacked even a clear understanding of which are these taxa. A recent study based on nine sites from eight countries across five continents was the first to identify surprising convergence of dominance in abundance and diversity (Srivathsan et al. 2023). Using a much larger dataset that spans more sites and includes an additional continent, we can here confirm nine of the ten families although switching quantification from diversity per trap to diversity per site. The only change is the inclusion of fungus gnats (Diptera: Mycetophilidae) which was previously ranked 15^th^. Notably, even the list of the top 20 taxa remains largely stable, with only two changes: Tachinidae and Agromyzidae (both Diptera) replace Bethylidae (Hymenoptera) and Sphaeroceridae (Diptera). Overall, we nevertheless find a striking global convergence of dominance across habitats and climates. This is encouraging in the sense that global biodiversity patterns can be uncovered by targeting largely the same taxa worldwide for ecology and biomonitoring: the ten dominant families identified here should become clear priorities for future biodiversity research.

Arguably, the global convergence in taxonomic dominance among flying insects was already unexpected, but another layer of surprise comes from our analysis of latitudinal diversity gradients. Of the top-ten dominant families, five show either no statistically significant gradient or an inverse one based on standardized samples from 27 countries. Only the remaining five families and the aggregate dataset follow a somewhat more expected pattern of increasing species richness toward the equator, although those increases are relatively weak compared to the data in the literature (Currie 1991; Blackburn and Gaston 1996; Hillebrand 2004; Kinlock et al. 2018). Note that these findings cannot be explained by lack of data. Indeed, we now have data for over 730,000 specimens in 66,505 species-level taxa for these top-ten taxa and the dataset for each one falls within the top quartile of dataset sizes used in previous meta-analyses of latitudinal diversity gradients (Fig. 2).

Latitudinal diversity gradients are among the most widely documented patterns in ecology, supported by decades of research across diverse taxa and spatial scales. Meta-analyses have shown that LDGs are generally evident whether diversity is measured locally (alpha) or regionally (gamma), suggesting a robust and scale-independent signal in biodiversity patterns (e.g., Willig et al. 2003; Hillebrand 2004; Kinlock et al. 2017). Our results challenge the generality of the LDGs for insect clades that dominate global species richness. For the top-ten taxa we find that alpha diversity does not consistently peak in the tropics. This could challenge explanations of LDG that act primarily on local species richness (Willig et al. 2003; Tuomisto 2010; Stein et al. 2014, such as those related to energy availability, climatic stability, or biotic interactions, in favor of mechanisms that influence species turnover. This rests on the assumption that beta diversity in these dark taxa will follow the classic latitudinal pattern, and indeed, preliminary evidence from a recent global-scale study (Seymour et al. 2024) hints at higher beta diversity in the tropics. To test this more directly, future studies should prioritize spatially structured sampling across closely spaced sites along latitudinal gradients, allowing for explicit evaluation of species turnover and its role in shaping regional richness patterns.

Encouragingly, the tools to carry out such fine-scale, transect-based studies are now available. Recent advances in high-throughput barcoding and imaging techniques allow large numbers of insect specimens to be efficiently assigned to putative species (DeWaard et al. 2019b; Srivathsan et al. 2019, 2021; Brydegaard et al. 2024; Meier et al. 2024). These methods make it possible to sample more evenly across taxa and geographic space, enabling a more comprehensive understanding of latitudinal diversity gradients. They also open the door to exploring how these patterns relate to body size. Previous studies have found a negative correlation between gradient strength and body size (Hillebrand and Azovsky 2001; Hillebrand 2004; Kinlock et al. 2018), and most species in the ten dominant flying insect clades examined here are relatively small. This may account for the weak gradients observed in some groups. However, it does not explain the pattern seen in Ichneumonidae, whose species tend to be larger than those in the other nine families.

Ichneumonidae not only represent the sole case of a significant inverse latitudinal gradient among our ten most diverse families, but also stand out as the clearest outlier relative to previous LDG studies. The effect size observed for the family ranks near the fifth percentile, placing it among the most extreme inverse gradients in our dataset. Indeed, the group has long been considered a textbook example of a taxon with an inverse latitudinal diversity gradient (Quicke 2015). However, the strength of the evidence had been questioned by recent tropical studies uncovering large numbers of undescribed species (Horstmann et al. 1999; Saaksjarvi et al. 2004; Veijalainen et al. 2012), and DNA barcoding data suggesting that tropical parasitoid communities may harbor high levels of cryptic diversity (Smith et al. 2006, 2008; Rodriguez et al. 2013). However, our results, based on standardized sampling across a broad geographic range, confirm a distinct inverse latitudinal gradient for Ichneumonidae and align with recent analyses of all sequence data for Ichneumonidae in the Barcode of Life Database (Castellanos-Labarcena et al. 2024). In our view, this long-standing discussion can now be considered settled for Malaise trap samples. Ichneumonid wasps show a markedly different latitudinal pattern from other dominant insect families, including the two other parasitoid top-ten taxa.

The global scale of our dataset enables, for the first time, a robust test of the two leading hypotheses proposed to explain the atypical diversity gradient in Ichneumonidae. The “resource fragmentation” hypothesis (Janzen and Pond 1975; Janzen 1981) suggests that tropical host biomass is split among too many rare, ecologically distinct species to support specialized parasitoids. The “nasty host” hypothesis posits that tropical herbivores accumulate more toxic compounds from their host plants, making them less suitable for parasitism (Gauld et al. 1992). Both hypotheses predict an increased proportion of generalist (idiobiont) species in the tropics, and earlier studies offered some support for this pattern (Gauld 1986, 1987; Hawkins et al. 1992). However, our data show no significant change in the proportion of idiobionts with latitude (Fig. 4d); if anything, the trend is slightly negative. Moreover, both idiobionts and koinobionts show similar declines in richness toward the tropics (Fig. S5). Since both hypotheses rely on ecological mechanisms that should affect all parasitoid groups, they also cannot explain the lack of a significant gradient in Braconidae and the positive gradient in Platygastridae. This suggests that explanations must lie beyond parasitoidism itself and may involve historical processes such as niche conservatism or phylogenetic inertia (Wiens and Donoghue 2004; Wiens and Graham 2005).

To conclude, our global analysis enables the confident identification of the top-ten clades of flying insects in Malaise trap samples, revealing a striking pattern of taxonomic convergence across continents, habitats, and climatic zones. At the same time, the expected strong latitudinal diversity gradients toward the tropics are lacking thus challenging long-held assumptions. The good news is that the top-ten families identified here are exceptionally well suited for comparative research: they are species rich, globally widespread, and sufficiently abundant to yield large data sets. This opens the door to for studies exploring which ecological or evolutionary factors best explain differences in diversity patterns. For instance, ecological hypotheses may be supported if top-ten taxa with similar gradients also share responses to environmental factors like precipitation or temperature seasonality. Conversely, evolutionary explanations may be better explanations if variation in gradient strength correlates with diversification history.

## Material and Methods

### Data mining and processing

We analyze DNA barcodes for 101 Malaise traps obtained at 45 sites in 27 countries, spanning all continents and biogeographic regions (Supplementary Material S1; Fig. 3). All but three of these datasets were retrieved from published studies (D’Souza et al. 2021; Chimeno et al. 2023; Srivathsan et al. 2023; Seymour et al. 2024); the three unpublished datasets were retrieved from the Barcode of Life Data System (BOLD) workbench, under the public campaigns folder “Global Malaise Program”. The 101 samples contained a widely varying number of specimens and our analyses are based on all sequences available for the same Malaise trap. Taxonomic assignments from the Barcode of Life Data System and Srivathsan et al. 2023 were accepted. This means that we continue to use a broad concept of Platygastridae that is equivalent to Platygastroidea that was recently formed by raising some clades to family rank (Chen et al. 2021).

### Sequence processing and objective clustering

To ensure that the datasets had sufficient overlap for multiple sequence alignments, short sequences were excluded: the length cutoff was 300 bp for the 313 bp barcode sequences (from Singapore) and 500 bp for the 658 bp barcodes (all others). Sequences that contained a stop codon when translated using the invertebrate mitochondrial genetic code were also excluded. Sequences were aligned using a modified version of transAlign.pl (Bininda-Emonds, 2005) upgraded to use Clustal Omega (Sievers et al., 2011) to perform a translation-based alignment. Objective clustering (Meier et al. 2006, https://github.com/asrivathsan/obj_cluster) was then used to group the specimens in clusters using a 3% distance threshold resulting in molecular operational taxonomic units (mOTUs), which were used as proxies for species diversity. For this study, “species” and mOTUs are used interchangeably through the text. Note that results of objective clustering have previously shown to be broadly congruent with those obtained with other distance-and tree-based approaches for species delimitation (Srivathsan et al. 2023). Sequence alignment and objective clustering analyses were initially performed for trap-level datasets individually which was used to identify the ten most diverse insect families in Malaise traps. Subsequently, for each of these ten families, family-level datasets were obtained by combining all specimens belonging to the same family from all 101 samples. We then used objective cluster to estimate the species diversity in each family-level dataset. For the 4 largest families, objective clustering could not be conducted directly due to the large size of the resulting pairwise matrix. Thus, only unique sequences were retained, and distances above 5% were excluded from the pairwise matrix. Objective clustering was conducted and specimen to cluster assignment was achieved by first adding the sequences that represented singletons at 5% distance and second by adding the specimen level information for each unique sequence. For all analysis, mOTUs corresponding to flying insect families only were used (Supplementary Material S2, Taxa removed).

### Community composition, abundance and diversity metrics

The top 10 most diverse families in Malaise traps were identified by ranking of average proportion of mOTUs per family (see Srivathsan et al., 2023). To avoid strong geographic biases deriving from some sites being represented by as many as 20 Malaise trap samples, we used the average values for the dominance and diversity metrics across all traps from a same site (defined as an individual reserve or national park, or city). To test if the overall catch of the Malaise trap samples conformed to the expected latitudinal gradient of more species, we generated a community matrix from the results of the objective clustering of each individual site. The matrix contained the abundance of each cluster per dataset; this allows the calculation of diversity metrics per site. We then calculated the diversity metrics of species richness and Shannon index (Magurran & McGill 2011) for each dataset using the *iNEXT* package (Hsieh et al. 2016) in R (R Core Development Team, 2021). Metrics were calculated using both size-based and coverage-based comparisons, using the average dataset size (13,503 specimens) and average coverage (0.8717) as a standardized sampling level, with sampling at specific datasets being extrapolated or rarefied as necessary to conform to that base level. The use of coverage instead of sample size for normalizing samples is, on theoretical grounds, preferable to rarefying datasets to equal sample size (Roswell et al. 2021). However, in our case, coverage levels varied widely across datasets, meaning several samples had to be extrapolated to over twice the reference sample size. In these conditions, the diversity metrics may be subject to a large prediction bias, and hence we report the results of both approaches, but mostly discuss the results obtained through size-level rarefaction.

For family-level datasets, a similar approach to trap-level datasets was applied: community matrices were generated and used to calculate diversity metrics. Datasets with fewer than 10 species were discarded to avoid biases caused by very low sampling and diversity metrics were calculated using the average number of specimens and average coverage levels for each family, and metrics for traps within the same size were averaged. We then fitted a linear regression model to investigate the relationship between diversity and latitude for each of the top 10 families.

To place the latitudinal diversity gradients observed in the top 10 families in a broader context, we compared our results to those of the datasets compiled in meta-analyses of latitudinal diversity gradient effect sizes (Hillebrand 2004; Kinlock et al. 2018). From the 970 individual latitudinal diversity gradient cases compiled in those studies, we retained only those that pertained to the terrestrial realm, covered at least 10 latitude degrees, and reported effect sizes for the latitudinal diversity gradient relationships. These criteria yielded 470 datasets. Note that only Kinlock et al. (2018) also reported the total number of species found in the study across its entire extent. The set of 114 studies used as an additional comparison point was drawn from all terrestrial studies in the latter meta-analysis that reported total species numbers and had >10 species.

To investigate the specific dynamics of abundance and diversity of Ichneumonidae when compared to the total insect communities, the proportion of specimens and mOTUs of Ichneumonidae within each dataset was calculated, again with values from different traps in the same site being averaged. Regression between relative abundance and latitude, and relative species richness and latitude were also calculated.

### Subfamily-level analyses in Ichneumonidae

To test whether the relative diversity of idiobiont (“generalist”) species of ichneumonids increased towards the tropics, we attempted to assign ichneumonid barcode sequences to a specific subfamily, as most subfamilies uniformly display either an idiobiont or koinobiont strategy (Broad et al. 2018). This assignment used BLAST (Basic Local Alignment Search Tool; Altschul et al. 1990) searches of our sequences against a library of ichneumonid sequences assembled as follows. First, we downloaded the full set of barcode sequences for Ichneumonidae available on BOLD (136,005 sequences in October 2023) in the form of a tab-separated text file containing sequence and metadata information. Of these, we only retained 91,225 sequences that were identified to genus level, with the assumption that specimens identified to genus level could be assigned reliable subfamily-level membership. Of these, 11,764 of the sequences corresponding to specimens included in our dataset were also removed, leaving 79,351 sequences that were used as the reference library.

We then ran BLAST searches for our full dataset of 21,354 ichneumonid sequences against the reference library under 80%, 85%, 90% and 95% cutoffs. Sequences that had a BLAST match under these specified cutoff were parsed into three groups: (1) “consistent”, if all top 10 BLAST hits matched consistently to same subfamily; (2) “multiple”, if the top 10 BLAST hits matched to multiple families; (3) “incorrect”, if one or more of the top 10 BLAST hits matched to a different subfamily than that present in the original ID available in the BOLD record. Only sequences in the “consistent” category were considered valid subfamily IDs based on BLAST and analyzed downstream. Also included in the analysis were query sequences that lacked a BLAST match at the specified cutoff but had a subfamily ID in the original sequence metadata. Overall, the 85% similarity cutoff allowed us to provide subfamily IDs for the largest proportion of species, with 16,394 sequences that had a “consistent” BLAST match and 2,530 sequences without a BLAST match that were previously identified to subfamily, for a total of 20,429 (95.7%) sequences with a final subfamily ID.

We assembled new community matrices for all taxa that could be assigned to a specific biological strategy via subfamily membership and calculated the diversity metrics for idiobiont and koinobiont taxa separately using the same parameters as described above, once again using only samples with at least ten ichneumonid species. We then fitted linear regressions between the observed diversity and the proportion of species belonging to each biological strategy against latitude.

Three of the subfamilies in our dataset cannot be entirely assigned to a single biological strategy. One is the small subfamily Adelognathinae, which represented only 0.52% of species in our overall assemblage. Two large subfamilies, Ichneumoninae and Pimplinae, have large numbers of both idiobiont and koinobiont species, and the specific biological strategy for each species could not be confidently assessed using only BLAST searches. Fortunately, the proportion of Ichneumoninae in the ichneumonid community was unchanged across latitudes (R^2^=0.01, *P*=0.498) so they are also unlikely to influence results. The Pimplinae, however, were proportionally more more common in the tropics (R^2^=0.20, P=0.002) and are predominantly idiobiont, so we performed a second set of subfamily-level analyses including the Pimplinae as part of the idiobiont group.

### Influence of bioclimatic variables

One of our goals was to test whether specific bioclimatic variables, rather than latitude per se, could be the subjacent factor explaining ichneumonid diversity patterns. Data on climatic conditions representing temperature, precipitation and seasonality were retrieved from the Worldclim database 1.4 (www.worldclim.org; Hijmans et al. 2005). We assessed multicollinearity among the predictor variables, comprising 18 bioclimatic variables obtained from WorldClim, using the Variance Inflation Factor (VIF). Predictor variables with VIF values > 10 were considered to exhibit strong multicollinearity and were excluded from the analysis (Kutner et al., 2004). For each of the top ten families, we generated a balanced model based on the Akaike Information Criterion (AIC) and calculated standardized coefficients to evaluate the relative effect sizes of the explanatory variables (Burnham and Anderson, 2002). The multi-model averaging and multicollinearity assessments were performed using the ‘MuMIn’ and ‘usdm’ R packages, respectively (Bartoń, 2023; Naimi et al., 2014). All analyses were conducted in the R statistical computing environment (R Core Team, 2023).

## Acknowledgments

The late Stefan Schmidt kindly granted early access to the data from the Chimeno et al. (2023) study. Nicole Kinlock provided information about the dataset and methods of her previous work. Gavin Broad and David Wahl contributed with advice on the prevalence of idiobionts and koinobionts among ichneumonid subfamilies.

## Author contributions

Conceived the study: B.F.S., R.M. Collected and edited data: B.F.S. Analyzed data: B.F.S., A.S., K.N. Wrote the manuscript: B.F.S., R.M., A.S. All authors contributed to draft revisions.

## Supplementary Materials

**Supplementary Material S1**. List of datasets of COI data used in this work. Each line corresponds to one individual Malaise trap operated for various time periods, including information about geographical coordinates, source of the data, and number of species and mOTUs for the complete dataset and each of the top 10 families.

**Supplementary Material S2**. Relative abundance and diversity data for 663 insect families for each of the 101 Malaise trap datasets. Comparison of top 10 most diverse taxa identified by Srivathsan et al. (2023) and this study including both the global dataset and specific geographic subsets.

**Supplementary Material S3**. Diversity metrics for the aggregate dataset and for each of the top 10 families, expressed both in species richness and Shannon index, standardized either by average number of specimens or average coverage across datasets. Sites with multiple Malaise trap samples are reported by the average across traps. Effect sizes for each of the top 10 taxa used in comparisons with published meta-analyses. Summary of spatial turnover data for the top 10 taxa.

**Supplementary Material S4**. Amount of variation in species richness explained by the multi-model (R^2^ and adjusted R^2^). Values of the relative importance of the variables included in the multi-model. Variable abbreviations: MTWQ = Mean Temperature Warmest Quarter; MTWetQ = Mean Temperature Wettest Quarter; TS = Temperature Seasonality; PWQ = Precipitation of Warmest Quarter; PWM = Precipitation of Wettest Month; PS = Precipitation Seasonality.

**Supplementary Material S5**. Metadata (effect size, slope, taxonomic richness, organism type, diversity type) for 470 terrestrial datasets included in the meta-analyses of Hillebrand (2004) and Kinlock et al. (2017), combined with our top 10 datasets.

**Supplementary Material S6**. Proportions and species richness by site for Ichneumonidae according to host use strategy (koinobiont versus idiobiont).

**Supplementary Material S7**. Species richness for Ichneumonidae by site with the removal of the largely temperate-restricted subfamilies Ctenopelmatinae and Diplazontinae.

**Supplementary Figure S1**: Jitter plots showing the proportion of species richness of the top 10 families among sites.

**Supplementary Figure S2**: Graph of effect size and slope of the regression line for the top 10 families datasets against a background of 470 terrestrial datasets in two broad meta-analyses investigating the strength of the latitudinal diversity gradient (Hilderbrand 2004; Kinlock et al. 2018). Colored dots correspond to the datasets for the top 10 families in this study. Strength of the latitudinal diversity gradient based on the Z-transformed correlation coefficient, and log slope calculated through a linear regression of estimated species richness and latitude.

**Supplementary Figure S3**: Linear regression of various diversity metrics against latitude in the aggregate Malaise trap dataset.

**Supplementary Figure S4**: Standardized regression coefficients, along with their 95% confidence intervals, to evaluate the influence of temperature and precipitation on the species richness of ten groups. Coefficients with 95% confidence intervals positioned entirely to the left or right of the dashed line were considered significantly different from zero. The symbol size reflects the relative importance of each variable, determined by summing the Akaike Information Criterion weights across all models in which the variable was included.

**Supplementary Figure S5**: Species richness of idiobiont (“generalist”) versus koinobiont (“specialist”) Ichneumonidae subfamilies by latitude.

