## Supplementary figures and images for "Weak and inverse latitudinal diversity gradients in the globally dominant flying insect clades"

### Supplementary Figure S1

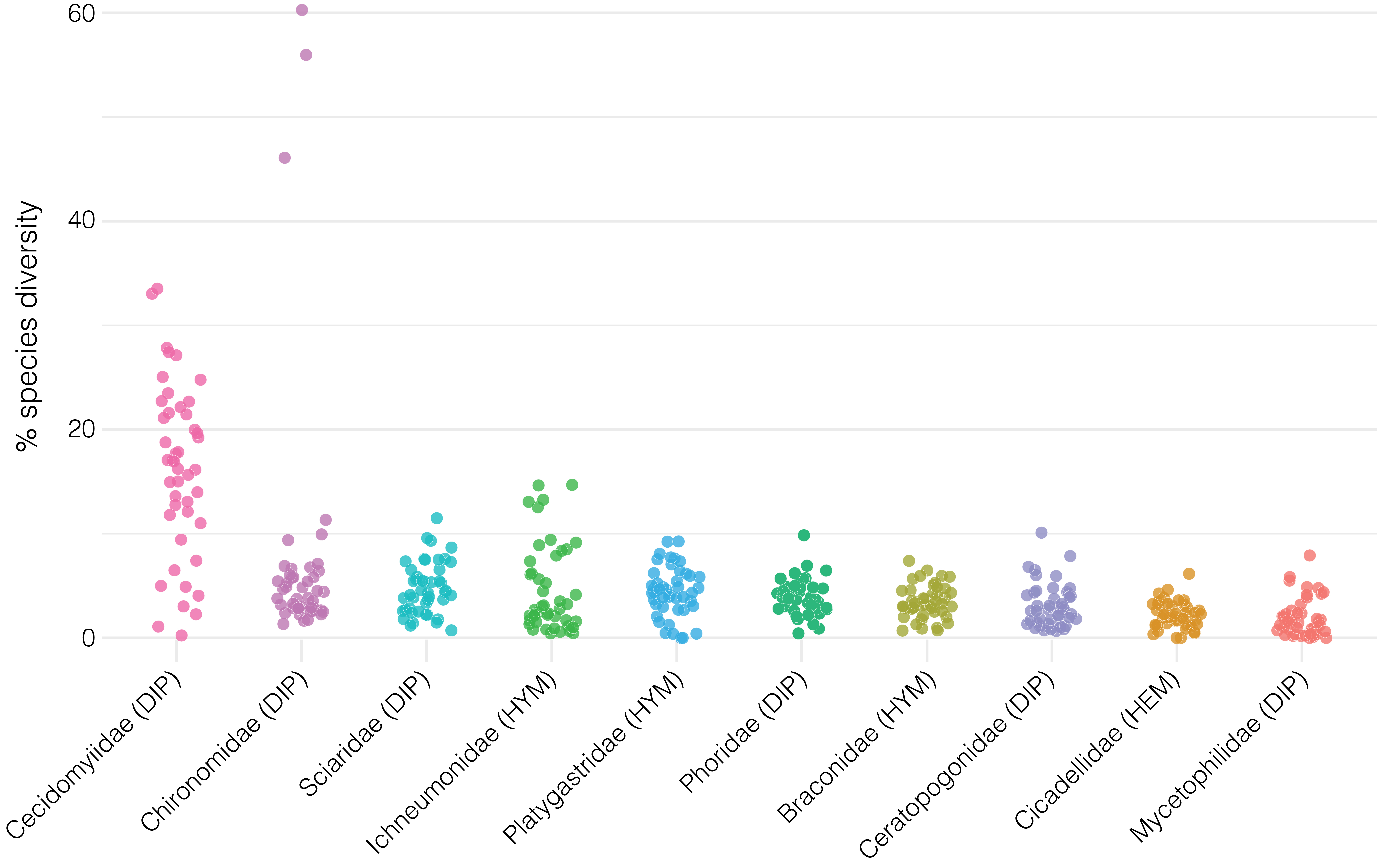

### Supplementary Figure S2

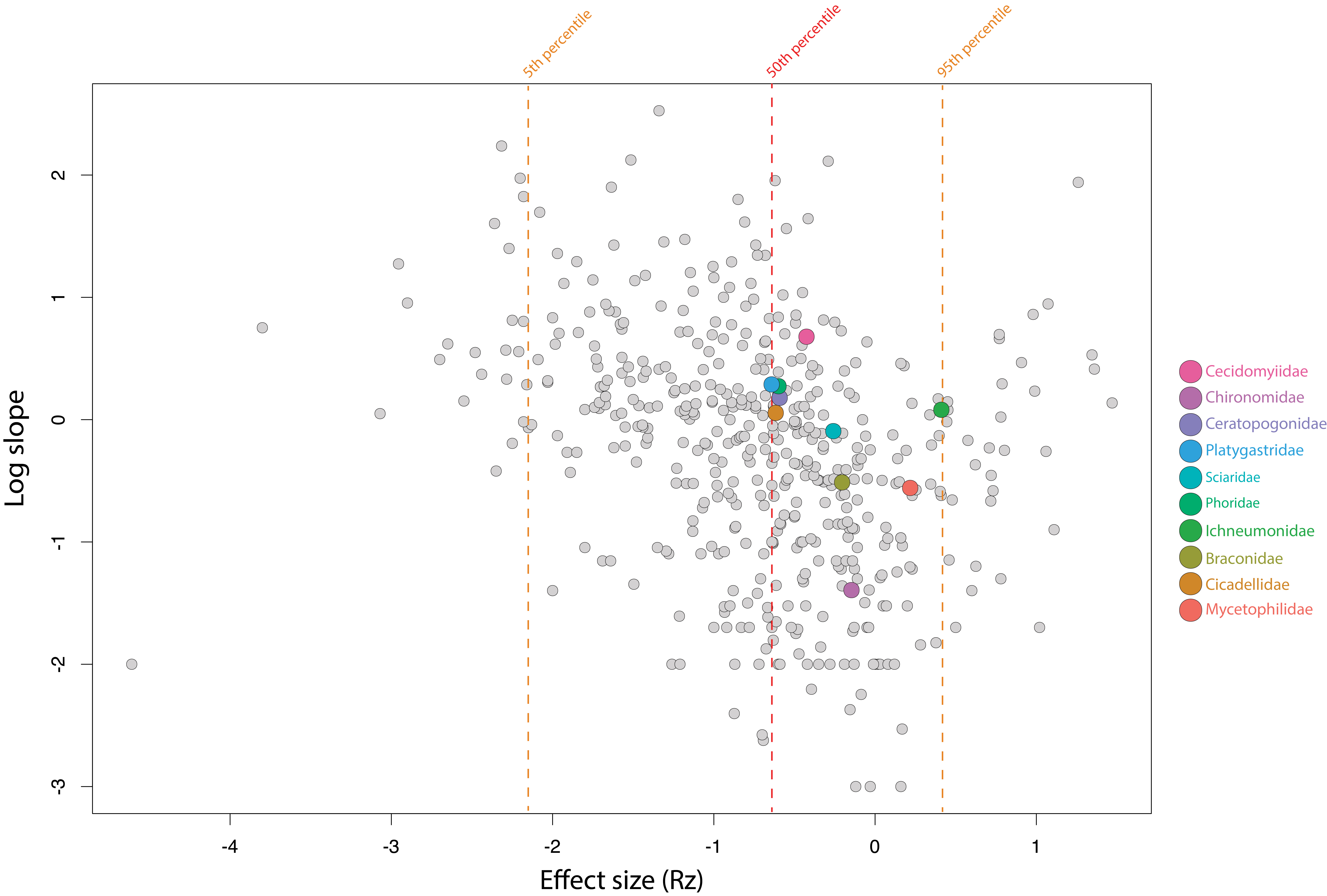

### Supplementary Figure S3

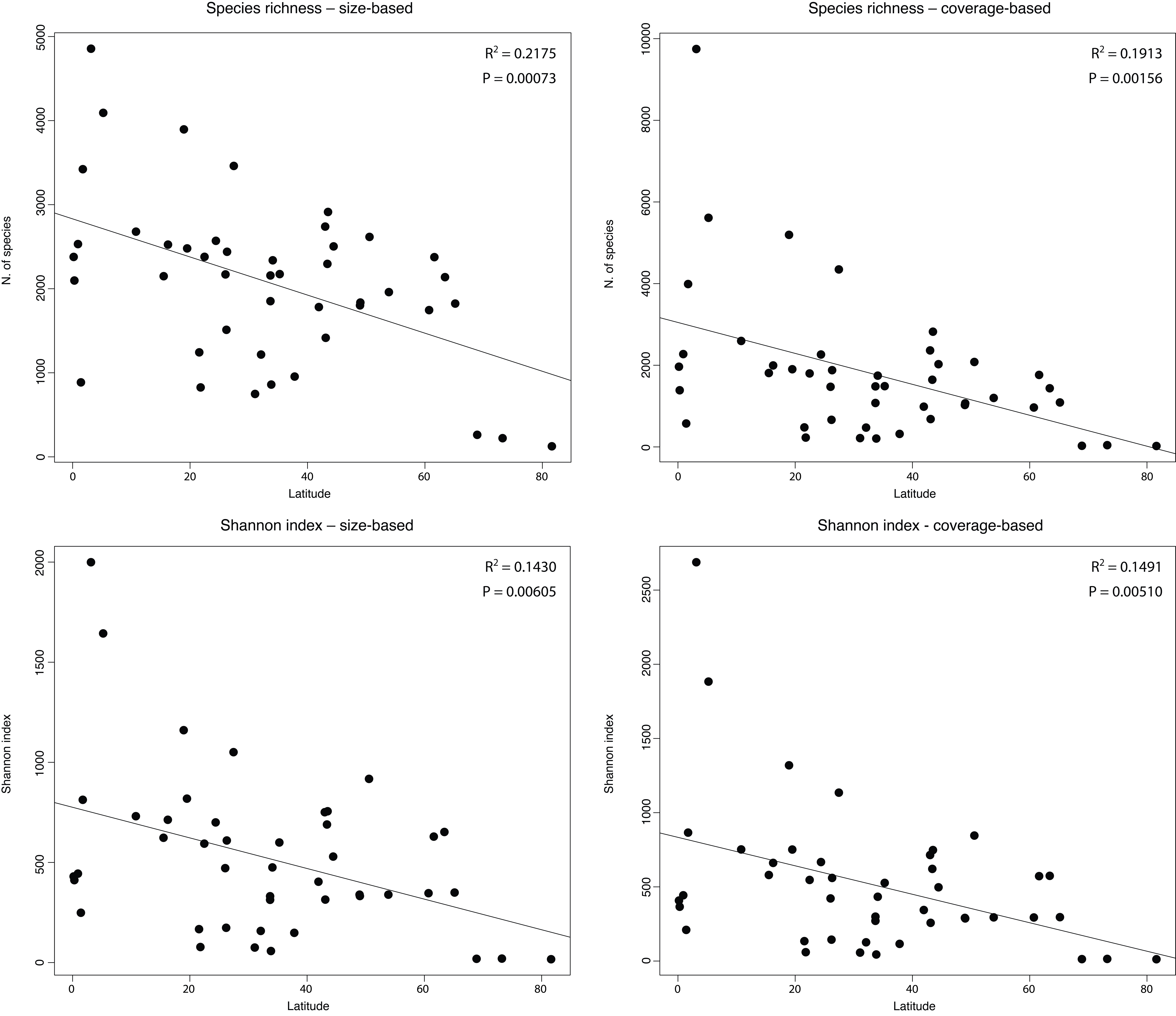

### Supplementary Figure S4

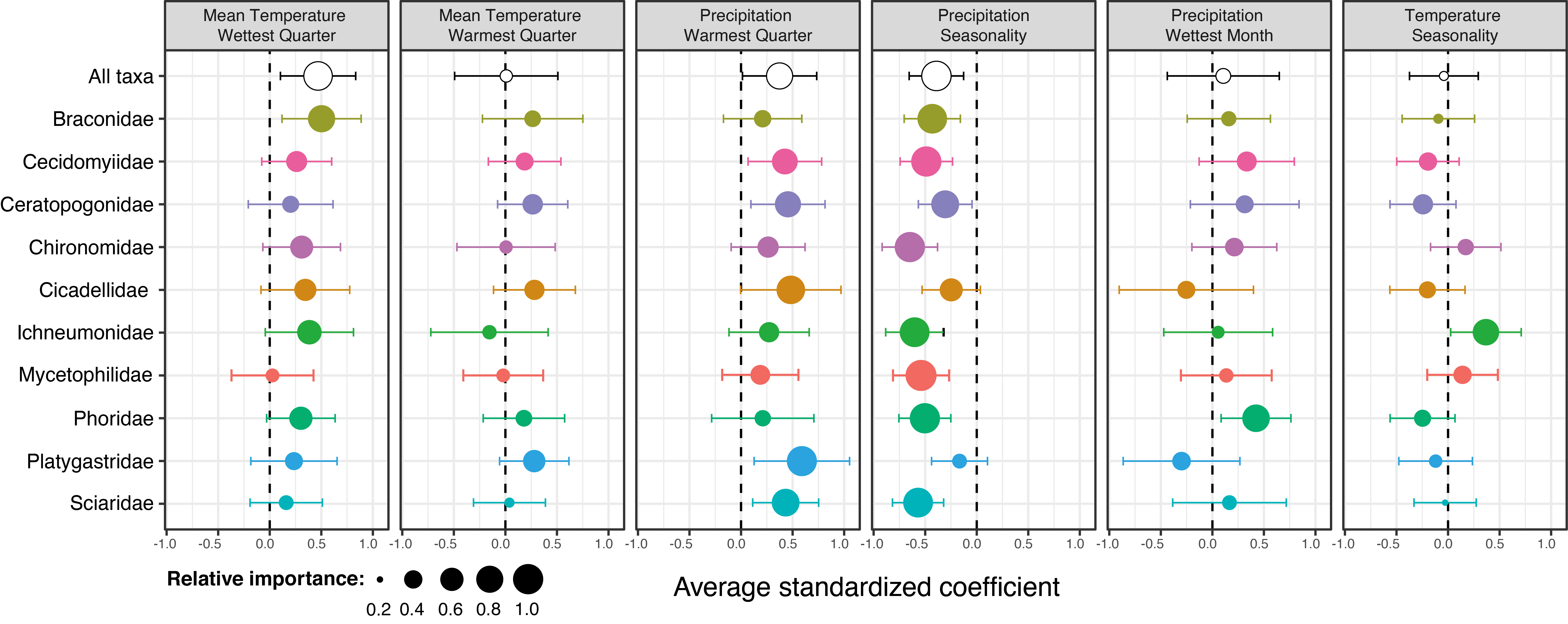

### Supplementary Figure S5

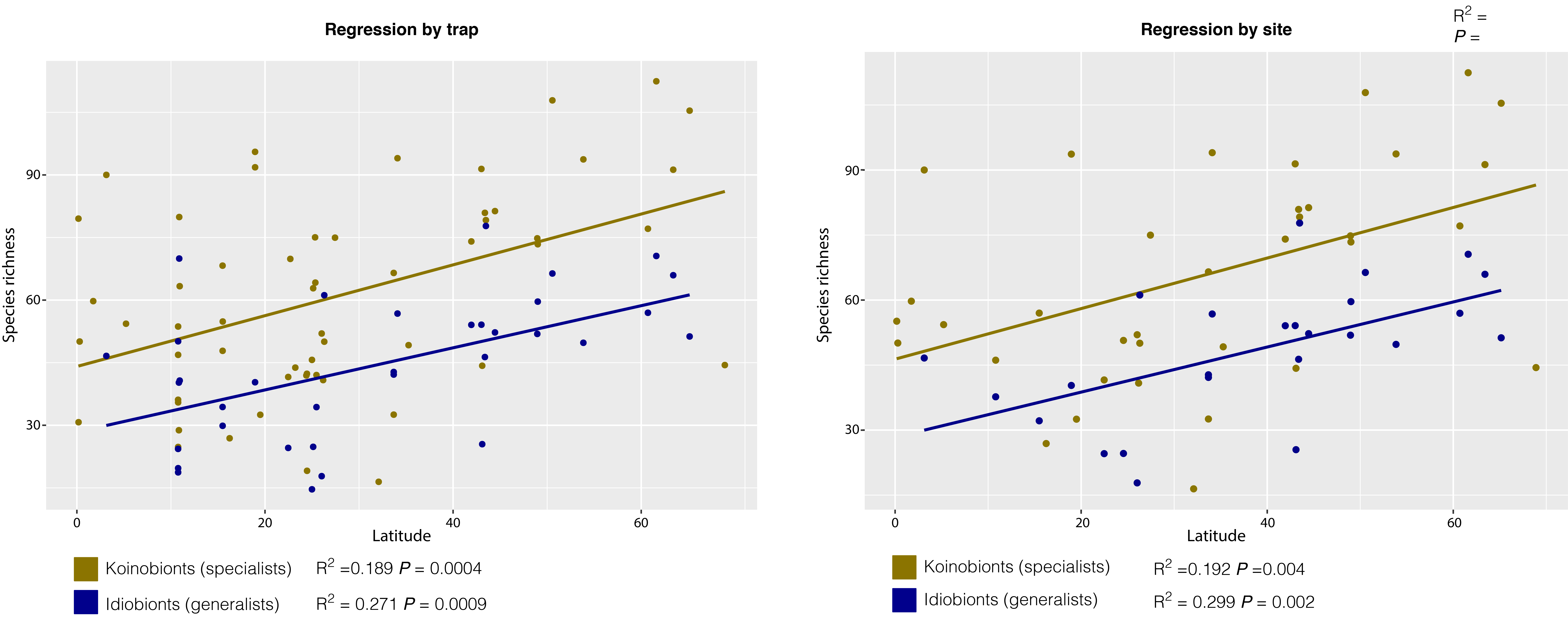
